# The bigger picture: global analysis of solubilization performance of classical detergents versus new synthetic polymers utilizing shotgun proteomics

**DOI:** 10.1101/2023.07.11.548597

**Authors:** Stefan Mueller, Jan Kubicek, Felipe Merino, Philipp Hanisch, Barbara Maertens, Jan-Wilm Lackmann

## Abstract

Integral membrane proteins are critical for many cellular functions. Roughly 25% of all human genes code for membrane proteins, and about 70% of all approved drugs target them. Despite their importance, laborious and harsh purification conditions often hinder their characterization. Traditionally, they are removed from the membrane using detergents, thereby taking the proteins out of their native environment, affecting their function. Recently, a variety of synthetic polymers have been introduced, which can extract membrane proteins together with their native lipids into a so-called native nanodisc. However, they usually show lesser solubilization capacity than detergents, and their general applicability for membrane protein biochemistry is poorly understood. Here, we used Hek293 cell membrane extracts and LC-MS-based proteomics to compare the ability of nanodisc-forming polymers against state-of-the-art detergents to solubilize the membrane proteome. Our data demonstrates the general ability of synthetic co-polymers to extract membrane proteins, rivaling the efficacy of commonly used detergents. Interestingly, each class of solubilization agent presents specific solubilization profiles. We found no correlation between efficiency and number of transmembrane domains, isoelectric point, or GRAVY score for any compound. Our data shows that these polymers are a versatile alternative to detergents for the biochemical and structural study of membrane proteins, functional proteomics, or as components of native lysis/solubilization buffers. Our work here represents the first attempt at a proteome-scale comparison of the efficacy of nanodisc-forming polymers. These data should serve as starting reference for researchers looking to purify membrane proteins in near native conditions.

## Introduction

Integral membrane proteins perform key cellular functions, such as cell signaling, selective transport, and more generally they mediate the interaction between cells and their environment. Although they represent about a quarter of all proteins encoded in the human genome (Fagerberg et al., 2010) they are the target of the majority of currently approved drugs (Santos et al., 2017). Despite their clear relevance, they remain very poorly characterized. For instance, up until 2021, only 3% of all structures in the PDB were membrane proteins (Le Bon et al., 2021). Many factors contribute to this, including low production yields, but perhaps the biggest challenge is that they need to be solubilized from the membrane. Traditionally, this is done using detergents, which remove the majority, if not all, lipids around the protein. Stripping membrane proteins of their native environment usually leads to instability as lipids are known to be important co-factors for protein structure and function (Barrera et al., 2013; Bogdanov et al., 2008; Contreras et al., 2011; Laganowsky et al., 2014; Pollock et al., 2018; Saliba et al., 2015; Schmidpeter et al., 2022), and detergents can replace the lipid bilayers only to a very limited extent (Seddon et al., 2004). In general, application and protein identity will determine which detergent is most appropriate. For example, quantitative and deep profiling of entire proteomes often requires either strong detergents such as sodium dodecyl sulfate (SDS), sodium deoxycholate (SDC), or RapiGest™, high concentrations of chaotropic agents such as 8 M Urea or 6 M Guanidinium Hydrochloride, or a mixture of them (Hughes et al., 2014; Vit et al., 2016; Wiśniewski et al., 2009). Furthermore, the enrichment of membranes by ultracentrifugation followed by washing with alkaline sodium carbonate or 8 M Urea has been described to improve the identification of low abundant membrane proteins (Fujiki et al., 1982; Kongpracha et al., 2022). In the case of quantitative proteomics, highly denaturing extraction conditions are extremely helpful, since they instantly reduce intrinsic protease and phosphatase activity in extracts, disrupt supramolecular protein networks and unfold proteins making them amenable for efficient tryptic digestion. In clear contrast, structural and functional characterization requires proteins in near native state, and enzymatic activity as well molecular interactions should be preserved as far as possible. While many soluble proteins can be easily extracted in simple aqueous buffers, the addition of mild detergents such as decyl maltoside (DM), dodecyl maltoside (DDM), and lauryl maltose neopentyl glycol (LMNG) (Choy et al., 2021) is required for membrane proteins. To stabilize extracted membrane proteins, detergents must be kept above their critical micelle concentration (CMC) over the entire workflow, significantly increasing reagent costs, experiment complexity, and negatively affecting many downstream applications.

More than two decades ago, the group of Jean-Luc Popot (Tribet et al., 1996) developed a new kind of amphipathic polymers or “amphipols”, capable of stabilizing integral membrane proteins. Amphipols can provide the hydrophobic environment the protein needs and bind it so tightly that no additional amphipol or detergent is needed in downstream applications. For this reason, classical amphipols like A8- 35, have been particularly popular for single particle electron cryo-microscopy (Cryo-EM) structures (Popot, 2018; Zoonens and Popot, 2014). Unfortunately, these polymers lack the ability to directly solubilize proteins from the membrane, and an initial detergent extraction is required. In 2009, the Overduin lab introduced a new class of co-polymer to tackle the challenge of solubilizing while preserving the native lipid environment (Knowles et al., 2009). Made by repeating units of styrene and maleic acid (SMA), SMAs were known to act as a molecular “cookie cutter” (Parmar et al., 2016). They extract integral membrane proteins together with a belt of their native lipids out of the membrane, forming a so-called Styrene and Maleic acid lipid particle (SMALP). To date, SMALPs have been used for many applications, including radioligand and spectroscopic binding assays, NMR based assays, and measurement of channel activity (see (Unger et al., 2021) for a summary). Unfortunately, SMALPs are unstable at pH values below 6.5, precipitate in the presence of divalent ions, and interfere with protein quantification using aromatic side chain absorption. However, in recent years, the promise of preserving the native lipid environment of the protein has led to the development of several alternative co-polymers that tackled some of the shortcomings of the original SMAs (for an overview of this rapidly evolving field see (Brown et al., 2021; Krishnarjuna and Ramamoorthy, 2022; Orekhov et al., 2022)). Variations in composition and manufacturing processes generate polymers with distinct capabilities. For instance, by replacing styrene with diisobutylene, Diisobutylene-maleic acid (DIBMA) produces a gentler polymer without absorption at 280 nm and higher tolerance to divalent ions (Oluwole et al., 2017). Although less efficient than SMAs, DIBMA could successfully solubilize ß2 adrenergic receptors producing a functional protein with higher stability compared to DDM (Harwood et al., 2021). Recently, two new polymer classes have been reported. Smith and colleagues (Smith et al., 2020) introduced a new co-polymers made of acrylic acid and styrene (AASTY) synthesized by reversible addition-fragmentation chain-transfer. AASTYs have a much more precisely controlled molecular weight and the use of acrylic acid confers them higher tolerance to divalent ions than SMAs (Timcenko et al., 2022). Interestingly, the group of Manuela Zoonens recently demonstrated that replacing the aliphatic group of traditional amphipols with cyclo- alkanes (CyclAPols, known commercially as Ultrasolute^TM^ Amphipols) resulted in polymers with excellent solubilization characteristics, capable of forming native nanodiscs (Marconnet et al., 2020). Moreover, CyclAPols solubilized bacteriorhodopsin shows increased thermal stability when compared to SMAs.

Lipidomics analysis of SMALPs and DIBMA solubilized lipid particles has demonstrated that both polymer classes can successfully extract native lipids from eukaryotic and bacterial cells, although the extracted lipidic composition is to some extend dependent on the exact chemical nature of the polymer. Interestingly, the lipid composition of SMALPs also depends on the identity of the protein (Teo et al., 2019) suggesting that these polymers can capture local compositional variations of the membrane. For most of these polymers, high-resolution Cryo-EM structures are available for membrane proteins solubilized from bacterial (Catalano et al., 2021; Doyle et al., 2022; Flegler et al., 2020; Higgins et al., 2021; Qiu et al., 2018; Sun et al., 2018; Swainsbury et al., 2023; Tascón et al., 2020) or eukaryotic cells (Yoder and Gouaux, 2020; Yu et al., 2021), many of which showing densities consistent with the presence of native lipids (Catalano et al., 2021; Doyle et al., 2022; Flegler et al., 2020; Qiu et al., 2018; Yoder and Gouaux, 2020). Unfortunately, while there is clear evidence that all these polymers can form native nanodiscs, most of the information about their solubilization capabilities comes from selected example proteins and systematic characterization of their solubilization capacity against other methods has only recently been performed for SMA (Carlson et al., 2019; Morrison et al., 2021).

Here, we used LC-MS analysis to compare the ability of nanodisc-forming polymers and detergents to extract membrane proteins from Hek293 cells. In contrast to the in-depth characterization of selected protein examples, shotgun proteomics facilitates the quantification of thousands of individual proteins in parallel and allows for more generalized observations. Testing a total of 22 extraction conditions including different types of SMAs, DIBMAs, AASTYs, and Ultrasolute^TM^ Amphipol, and the commonly used non-ionic detergents DM, DDM, and LMNG, our data represents the most comprehensive proteome-scale study on membrane protein solubilization by polymers.

## Methods

### Cell culture

Human embryonic kidney 293 cells (Expi293F™, Thermo Fisher) were cultured in Expi293™ Expression medium at a cell density between 5 x 10^5^ cells/ml and 4 x 10^6^ cells/ml in suspension. 100 ml of a 2 x 10^6^ cells/ ml culture growing in exponential phase were harvested by centrifugation and the cell pellet was stored at – 80 °C.

### Preparation of membrane pellets

The cell pellet was resuspended in 150 mM NaCl and 20 mM HEPES, pH 7.4, cells were disrupted by sonication. After a centrifugation step at 9,000 x g the protein concentration of the supernatant was adjusted to 0.25 mg/ml using 150 mM NaCl and 20 mM HEPES pH 7.4. The adjusted supernatant was pipetted into 100µl aliquots, and the aliquots centrifuged at 60,000 x g for 1 hour. The supernatant was discarded and the cell pellets consisting of approximately 25 µg of total proteins were frozen at -80 °C.

### Protein extraction and digestion

For an initial protein extraction screen SDS was tested as a 5% solution in 50 mM HEPES pH 7.4. RIPA buffer was composed of 150 mM NaCl, 1% Nonidet P-40, 0.5% deoxycholate, and 0.1% SDS in 50 mM HEPES pH 7.4. All other tested synthetic polymers and detergents were used as 2.5% solutions in 50 mM HEPES pH 7.4 (Table 1). In a subsequent dilution experiment the extraction properties of a selected panel of detergents and polymers were tested using five different concentrations ranging from 0.125% to 2.5% in 50 mM HEPES pH 7.4 (Table 1). All reagent solutions and samples were used at 5° C and kept in pre-chilled cooling racks. Either 25 µl detergent or polymer solution were added to the membrane pellets, each containing approximately 25 µg total protein. The tubes were briefly vortexed and placed in a Thermomixer (Eppendorf) for 16 h at 800 rpm and 5°C. After extraction, the tubes were transferred to a pre-chilled rotor and centrifuged for 35 min at 21,000 x g and 5°C. 20 µl supernatant was carefully transferred to new tubes without touching their bottom. 80 µl of cold acetone (-20°C) was added, the tubes were briefly vortexed and incubated for 120 min at -20°C to precipitate proteins. Supernatants were removed and the remaining protein pellets were washed once with 250 µl cold acetone (-20°). 25 µl 2% deoxycholate in 50 mM HEPES Tris HCl pH 8.5, 5 mM TCEP, and 40 mM chloroacetamide were added to the airdried protein pellets. Proteins were re-solubilized, reduced and S-alkylated in a single step for 2 h at 46°C. 0.5 µg trypsin was added in 5 µl 2% deoxycholate in 50 mM TrisHCl pH 8.5 and proteins were digested at 37°C. After 16 h, 30 µl of 5% trifluoroacetic acid was added to stop the digestion. By following this precipitation-resolubilization protocol, we removed solubilization agents and standardize the tryptic digestion step, which is fully compatible with SDC (Lin et al., 2008). Samples were centrifuged for 5 min at 21,000 x g and 20°C to pellet precipitated deoxycholate and 50 µl clear supernatant was removed and transferred to fresh tubes. 20 µl were subsequently loaded on DVB-RPS mixed mode stage tips activated with 20 µl methanol and equilibrated to 0.1% formic acid (FA). The loaded peptides were sequentially washed with 40 µl 0.1% FA in water and 40 µl 0.1% FA in 60% acetonitrile followed by alkaline elution with 40 µl 1% ammonium hydroxide in 60% acetonitrile. Eluates were immediately acidified with 10 µl 10% FA and dried in a vacuum centrifuge. Samples were re-solubilized in 10 µl 5% FA, 2% acetonitrile, and stored at -20°C until LC-MS analysis.

**Table 1.** Summary of the solubilization agents used in our screening experiments.

| Class | Copolymer/detergent | Primary screen | Dilution screen |
| --- | --- | --- | --- |
|  |  | 2.5% | 0.125 – 2.5% |
| AASTY | AASTY 11-55 | yes | No |
| AASTY | AASTY 6-55 | yes | Yes |
| AASTY | AASTY 11-50 | yes | No |
| AASTY | AASTY 6-50 | yes | Yes |
| AASTY | AASTY 11-45 | yes | Yes |
| Amphipol | Ultrasolute™ Amphipol 18 (CyclAPol C8C0-50) | yes | Yes |
| Amphipol | Ultrasolute™ Amphipol 17 (CyclAPol C6C2-50) | yes | Yes |
| DIBMA | DIBMA Glucosamine | yes | No |
| DIBMA | DIBMA Glycerol | yes | Yes |
| DIBMA | DIBMA 12 | yes | Yes |
| DIBMA | DIBMA 10 | yes | Yes |
| SMA | SMA 502-E | yes | Yes |
| SMA | SMA 300 | yes | Yes |
| SMA | SMA 200 | yes | Yes |
| Neutral detergents | Digitonin | yes | Yes |
| Neutral detergents | LMNG | yes | Yes |
| Neutral detergents | DM | yes | No |
| Neutral detergents | DDM | yes | Yes |
| Ionic detergents | RIPA buffer (1 % Nonidet P-40, 0.5 % deoxycholate, 0.1 % SDS) | yes | No |
| Ionic detergents | SDS | yes, 5% | No |
| Ionic detergents | SDC | yes | No |
| Ionic detergents | Fos-12 | yes | No |
| Negative Control | HEPES | yes | Yes |

### LC-MS Data acquisition and processing

Protein digests were analyzed on a Q-Exactive Plus (Thermo Scientific) mass spectrometer that was coupled to an EASY nLC 1200 UPLC (Thermo Scientific). Re-solubilized protein digests were loaded onto an in-house-made pulled tip column (30 cm length, 75 µm inner diameter, filled with 2.7 µm Poroshell EC120 C18, Agilent) which has been equilibrated in 0.1% FA in water. Peptides were separated using increasing acetonitrile concentrations in 0.1% FA followed by a 10 min column wash with 95% acetonitrile in 0.1 % FA. 140 min gradients were used for data dependent (DDA) acquisitions and 90 min gradients with data independent (DIA) acquisitions. The protein extraction screen experiment was acquired with the DDA method. Briefly, MS1 survey scans were acquired from 300 to 1,750 m/z at a resolution of 70,000 and stored in profile mode. The top 10 most abundant peptides were isolated within a 1.8 Th window and subjected to HCD fragmentation at a normalized collision energy setting of 27%. The AGC target was set to 5e5 charges, allowing a maximum injection time of 60 ms. Product ions were detected in the Orbitrap at a resolution of 17,500. Precursors were dynamically excluded for 25 s. A DIA method was used for the analysis of the dilution experiment. MS1 scans were acquired from 380 m/z to 1,020 m/z at 35,000 resolution. Maximum injection time was set to 60 ms and the AGC target to 1E6. MS2 scans starting from 250 m/z were acquired with 17,500 resolution, maximum injection time of 60 ms, and the AGC target set to 1E6. 30 x 20 m/z staggered DIA windows covered the precursor mass range from 400 m/z – 1,000 m/z and resulted in 60 nominal 10 m/z windows after deconvolution. For spectral library generation, aliquots of the 2.5% condition of all extraction experiments were pooled and analyzed with six consecutive gas phase fractionation (GPF) runs covering the mass range from 400 m/z to 1,000 m/z in sequential 100 m/z segments. MS2 scans were acquired with 25 x 4 m/z staggered windows in a 100 m/z range. All other settings were identical to the standard DIA method used for sample acquisition. DDA and DIA data were stored as centroid.

## Data analysis

DDA runs were processed with MaxQuant version 1.5.3.8 (Tyanova et al., 2016a) using default parameters. Briefly, MS2 spectra were searched against the human canonical Swissprot reference proteome database (UP5640, downloaded 08/26/2020), including a list of common contaminants. False discovery rates on protein and PSM level were estimated by the target-decoy approach to 1% Protein FDR and 1% PSM FDR respectively. The minimal peptide length was set to 7 amino acids and carbamidomethylation at cysteine residues was considered as a fixed modification. Oxidation (M) and Acetyl (Protein N-term) were included as variable modifications. The “match between runs” option was enabled within replicate groups. Protein and gene identifiers together with iBAQ intensities were imported into Perseus version 1.6.5.0 (Tyanova et al., 2016b) for subsequent analysis. Decoys and potential contaminants were removed. iBAQ values were log2 transformed and identified proteins were filtered for at least 3 out of 3 values in at least one replicate group. The number of transmembrane domains (TMDs) together with start and end points of corresponding TMD sequence stretches were extracted from Uniprot for each protein and merged to the data set. Principal component analysis was calculated on the imputed group means using Perseus defaults. Hek293 gene expression data were downloaded from https://www.proteinatlas.org (Uhlén et al., 2015). Entries with transcripts per million (TPM) below 1 were considered to be noise level and removed together with non-membrane proteins (TMD count = 0). Remaining entries were considered as expected membrane protein expression background. Grouped mean iBAQ intensities as well as protein level TMD counts were merged to the expression background and the percentage of entries that were also identified by the proteomics experiment was calculated to estimate the analytical depth of the proteomic analysis. Gravy Scores and Isoelectric points were calculated from the human canonical Swissprot database proteome (UP5640) using the “Peptides” R-package (https://github.com/dosorio/Peptides/).

All DIA raw files were deconvoluted and transformed to mzML files using the msconvert module in Proteowizard. DIA-NN 1.8.0 (Demichev et al., 2020) was used in library free mode to generate a project specific spectral library from a human canonical Swissprot reference proteome fasta file and the six GPF runs. The settings differing from defaults were: min precursor m/z set to 400, max precursor m/z set to 1,000, min fragment m/z set to 250, max fragment m/z set to 1,800, and precursor charge states ranging from 2 to 4. The resulting spectral library (11,214 proteins, 11,186 genes and 139,723 precursors) was used for the subsequent analysis of the sample runs with default parameters and the relaxed protein inference option. Using the “diann” R-package (https://github.com/vdemichev/diann-rpackage). MaxLFQ intensities were calculated from non-normalized intensities provided in the DIA-NN output file and were imported into Perseus. For heatmap generation, log2 transformed MaxLFQ values were normalized to an interval from 0-1.

## Results and Discussion

We used LC-MS to compare a panel of nanodisc-forming polymers and commonly used detergents for their ability to extract integral transmembrane proteins from Hek293 cell membranes. In total 22 extraction conditions including different types of SMAs, DIBMAs, AASTYs, and Ultrasolute^TM^ Amphipols, and the commonly used non-ionic detergents DM, DDM, and LMNG were tested. Our non-detergent control was HEPES buffer. As positive control we chose SDS, SDC, and Fos-12 as a panel of very aggressive detergents, as well as the ubiquitously used RIPA buffer. While these are not suited for native extraction, their high protein solubilization capacity should provide an upper limit of detectable membrane proteins in the current experimental setup. For an overview of the compounds tested (Table 1) and their physicochemical characteristics see Table S1.

### Analytical depth of our study

We could identify a total of 7,197 proteins across all runs, including 5,665 non-membrane proteins (Table S2). We attribute this to loosely membrane-interacting proteins or soluble proteins trapped in vesicles. Normally, this is dealt with by washing the samples with alkaline carbonate buffers (Fujiki et al., 1982; Kongpracha et al., 2022). However, the elevated pH needed for this (> 11) is incompatible with native protein extraction. Therefore, we filtered the data for proteins with at least one trans membrane domain (TMD), leaving 1,532 membrane protein identifications. For comparison, RNA-seq experiments of Hek293 cells report 2,371 transmembrane proteins to be expressed with transcript levels greater than 1 transcript per million (Protein Atlas; https://www.proteinatlas.org/). Considering this number as expression background, we estimated the analytical depth of our proteome analysis to be better than 64 % for proteins with at least one TMD. Importantly, while mRNA levels are not always good predictors of protein amounts, they should be strongly correlated with the cell growth conditions we tested (Liu et al., 2016). Indeed, comparing transcript levels with our detected proteins (Figure S1), suggests that low gene expression is the major reason for proteins missing in our data set, strongly arguing for the high quality of the dataset.

### Different solubilization compounds result in unique solubilization patterns

For our initial screening round, we assessed the solubilization capacity of all compounds at a final concentration of 2.5%, or 5% in the case of SDS. To get an initial overview of their solubilization differences, we used the mean abundance of the detected membrane proteins to perform a principal component analysis (PCA). Strikingly, the first two components (∼70% of the variance) reveal that individual reagents form tight clusters with other members of their respective classes (Figure 1A). Higher components can separate ionic from non-ionic detergents, while to a lesser extent still preserving the separate clustering of the different polymer classes (Figure 1B)

**Figure 1.**
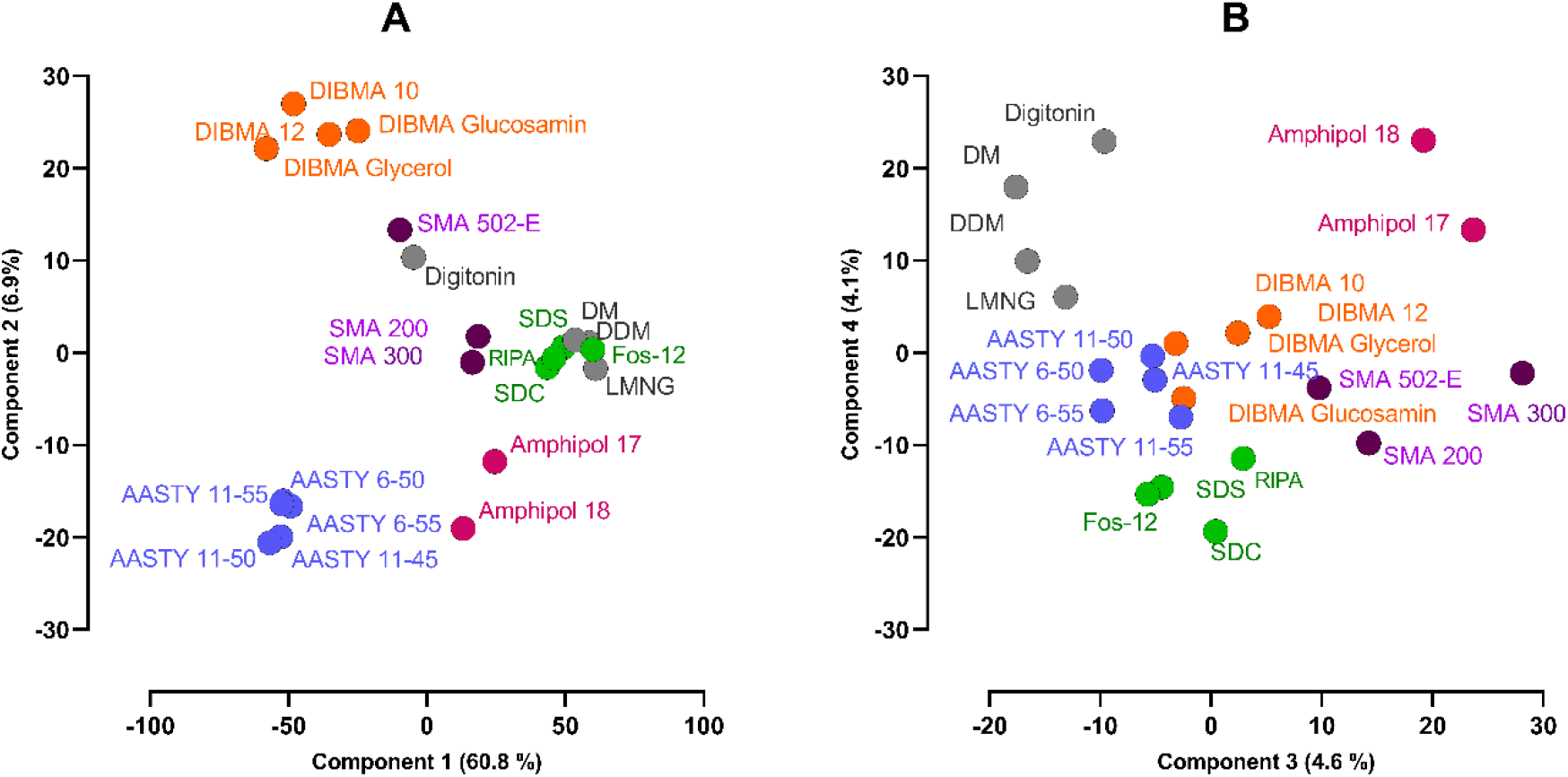
Clustering of membrane protein extraction profiles of different solubilization agents. Principal component analysis of the membrane protein iBAQ values. **A.** Components 1 and 2 of the PCA. **B.** Components 3 and 4. The percentage in each axis label shows the proportinal size of the corresponding eigenvalues. The means of three replicates per extraction condition were used. Each class of solubilization agent is shown in a different color to highlight their clustering.

The clear separation between detergents and polymers, as well as the separation between different polymer classes suggest that each solubilization agent class produces a distinctive fingerprint. However, a relation between the individual components of the PCA and the specific physicochemical characteristics of the compounds is not immediately obvious. To our surprise, although denaturation potential and extraction properties of ionic and non-ionic detergents are clearly different, DM, DDM, and LMNG cluster closely with the much more aggressive SDS, SDC, and Fos-12. As expected, the RIPA buffer — a mixture of SDS, SDC, and Nonidet P-40 (NP-40) — is located within this cluster as well. Digitonin does not follow this trend and is located far away from the main detergent cluster, near SMA 502-E.

### Polymers can efficiently solubilize membrane proteins

Next, we sought to understand whether the differences in solubilization profiles we observe are a consequence of differences in solubilization efficiency or versatility. In our negative control, we detected an average of 253 membrane proteins, 181 (71.41%) of which are single span membrane proteins (Figure 2). In clear contrast, the membrane protein counts for the ionic detergents SDC, SDS, and Fos-12 were 1250, 1,241, and 1,280 respectively. Unexpectedly, the ionic detergents performed similarly to Ultrasolute^TM^ Amphipol-17 and Amphipol-18 (1,270 and 1,272 membrane proteins respectively) and were outperformed by the non-ionic detergents DM, DDM, and LMNG (membrane protein counts of 1,297, 1,308, and 1,315 respectively); the latter being the highest in the data set (Figure 2). We obtained intermediate efficiency for SMAs, with counts of 1,175, 1,198, and 1,066 for SMA-200, SMA-300, and SMA-502-E respectively. AASTYs and the DIBMAs had the lowest performance with counts ranging from 991 to 1,010 and from 897 to 969 respectively (Figure 2). Nevertheless, DIBMAs and AASTYs are efficient solubilization agents, showing significantly higher solubilization than the background HEPES alone. Importantly, solubilization efficiency of membrane proteins is only one piece of the puzzle. Purification strategy, stabilization of the membrane protein, and downstream applications are equally important aspects to consider, which makes screening of various reagents indispensable. Indeed, polymer identity or even concentration can affect all of these parameters (Szundi et al., 2021). Absolute solubilization efficiency may also not be critical for many applications where a near-native state is the strongest constraint and sufficient initial material is present; the obvious example would be after overexpression. For instance, the comparably milder DIBMA has been successfully used to determine the Cryo-EM structure of the mechanosensitive-like channel YnaI (Flegler et al., 2020) and has been successfully used to produce highly active and stable GPCR samples (Harwood et al., 2021).

**Figure 2.**
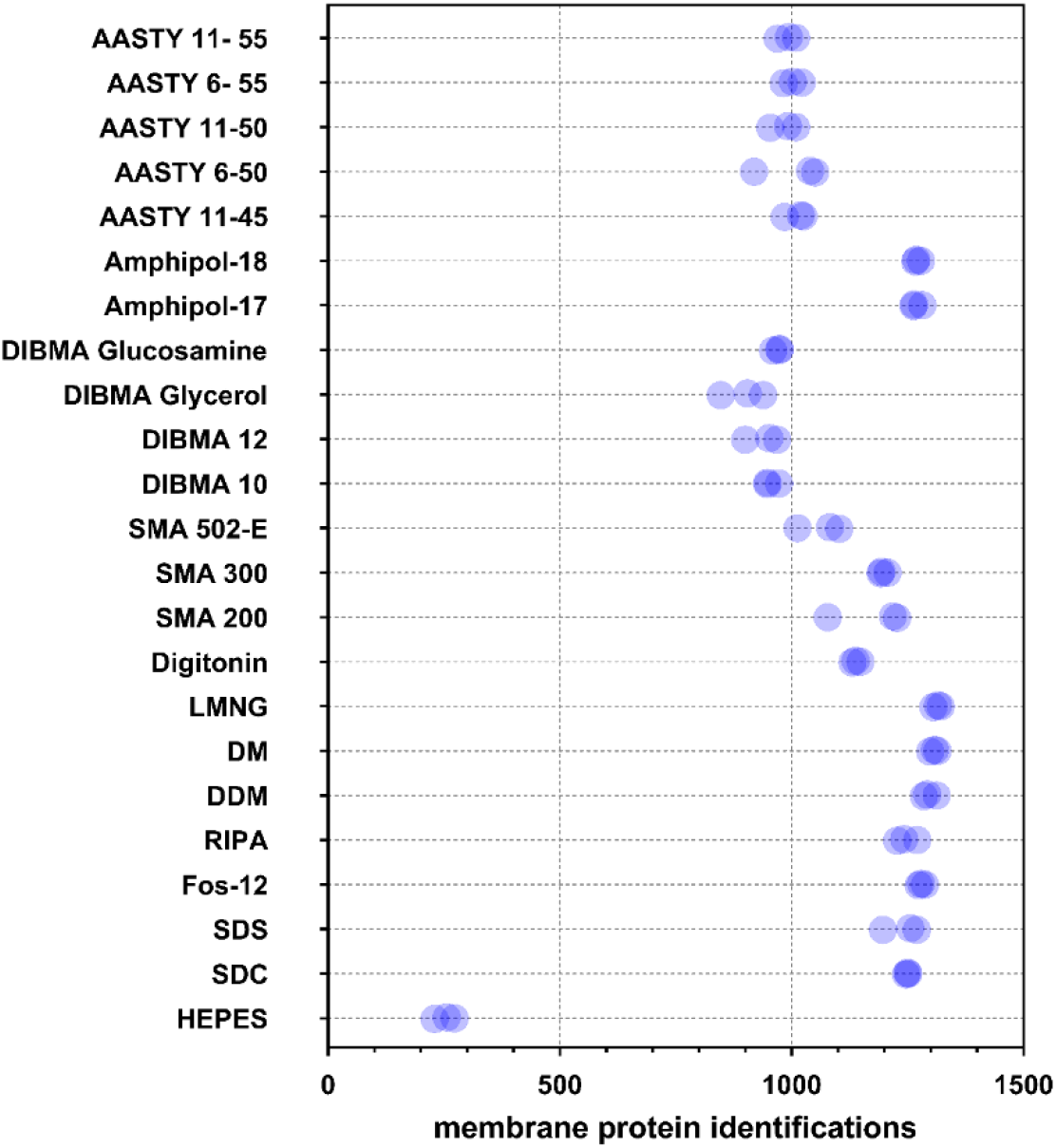
Cumulative number of detected membrane proteins. For each replicate and condition, the sum of membrane protein identifications is shown.

### Polymers can efficiently extract diverse classes of membrane proteins

Our PCA analysis revealed that the different compound classes we tested produce distinct solubilization patterns. Although they clearly differ in their solubilization efficiency, this cannot be the only driver as the clustering of detergent and polymer classes remains present for higher components in the PCA analysis (Figure 1B). We first analyzed the potential impact of the membrane protein topology on solubilization. For this, we grouped proteins according to their number of TMDs and calculated identification rates as the fraction of membrane proteins we identified versus what we expected from RNA data. Identification rate differs substantially among TMD groups (Figure 3). Interestingly, the profiles are similar for all tested extraction conditions, with scale differences clearly proportional to efficiency. Since median protein abundancies have a major impact on identification rates in proteomic experiments, we tested if this was behind the identification rate pattern. Indeed, the mRNA level shows a similar pattern to our detection rates (Figure 3), with a significant correlation between identification rates and median mRNA abundance (Figure S1). In turn, this suggests that protein abundance and not preferential solubilization is behind the identification rate pattern across the TMD groups.

**Figure 3.**
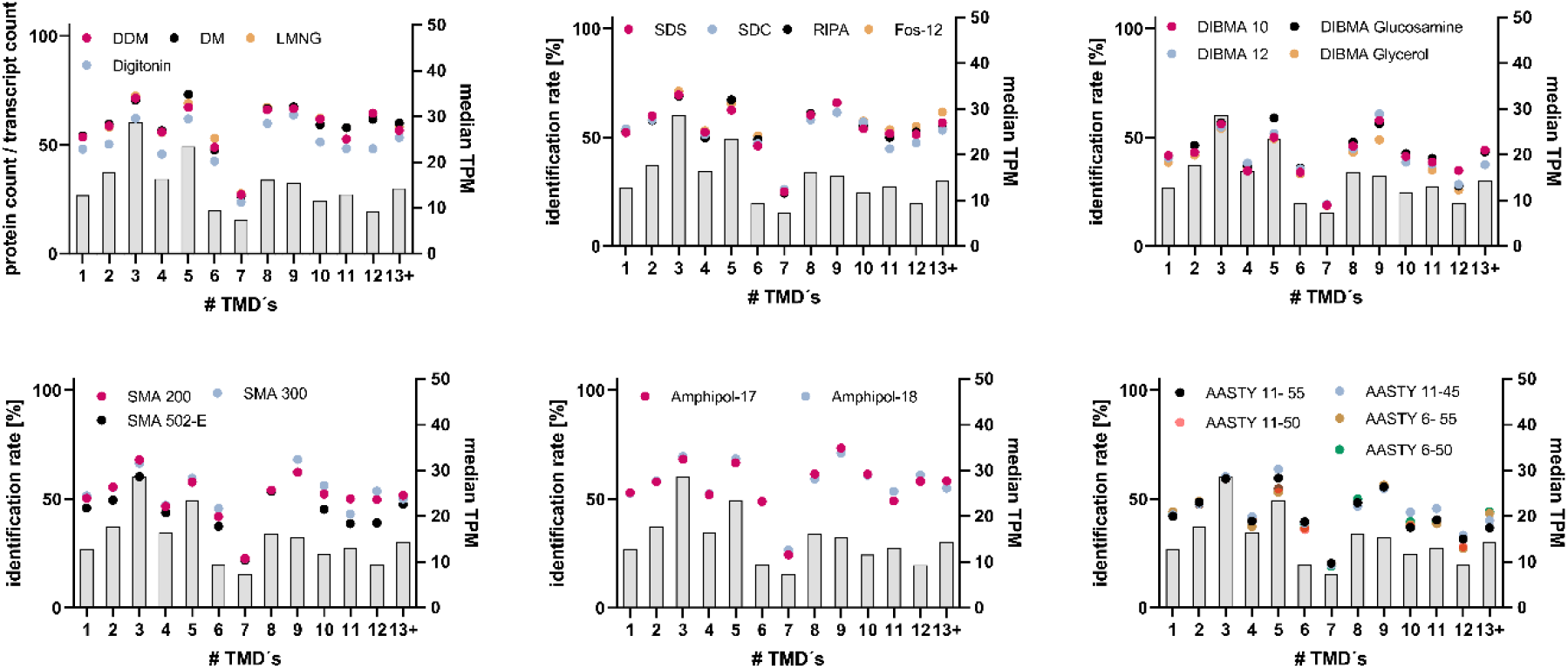
Membrane protein identification rate follows protein abundance in relation to number of TMDs. Proteins were grouped according to their number of transmembrane domains. For each solubilization agent class and each group, the identification rates (colored dots) were calculated as fraction of identified membrane proteins versus the total number of expressed membrane protein transcripts in Hek293 cells in the respective group. For reference, the RNA levels were plotted as median number of TMPs in grey bars.

A clear candidate for the differential polymer preferences would be the physicochemical properties of the proteins and their respective transmembrane regions. For instance, hydrophobicity and charge substantially differ across membrane proteins and could easily affect the polymer’s solubilization efficiency. To test this, we calculated GRAVY scores and isoelectric points for either the full protein sequences or their membrane spanning regions. Interestingly, the distributions of either characteristic for the detected membrane proteins reveal no major difference among the solubilization agents, strongly suggesting that isoelectric points and hydrophobicity cannot explain the differential extraction patterns that we observe (Figure 4).

**Figure 4.**
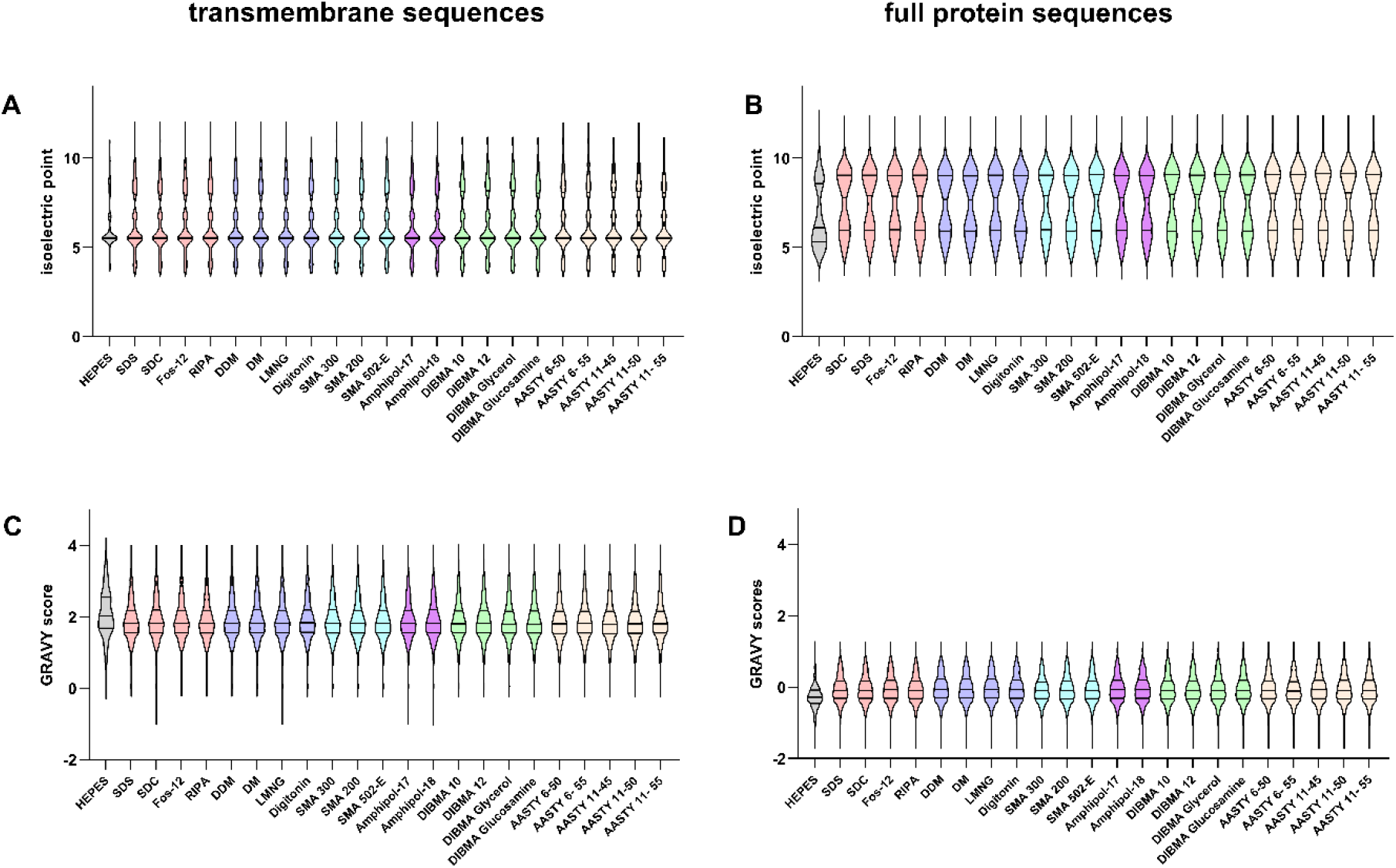
Distribution of isoelectric points and GRAVY scores for the proteins detected in solubilization experiments. For all detected membrane proteins, isoelectric points (**A,B**) and GRAVY scores (**C,D**) were calculated using either the transmembrane regions (**A,C**) or the full protein sequence (**B,D**).

Contrary to detergents, the polymers we tested solubilize the protein together with its lipidic microenvironment. Far from being just a scaffold for these proteins, the exact lipidic environment around them is matched to their specific requirements (Lee, 2004), with each of them possessing its own lipidic fingerprint (Corradi et al., 2018). Interestingly, SMAs have already been shown to solubilize, from the same membrane, proteins with different local lipidic compositions (Teo et al., 2019). However, early characterization of nanodisc-forming polymers has shown that their solubilization efficiency is somehow dependent on membrane composition (Oluwole et al., 2017; Smith et al., 2020). Combined, this suggests that, for a specific target, differences in solubilization ability between polymers would be partially determined by the identity of the annular lipids surrounding the membrane protein.

### Optimal solubilization conditions are specific to each compound and target combination

So far, we compared the capacity of polymers and detergents to extract membrane proteins using a standard concentration of 2.5%. Although this concentration is effective for all conditions tested, it is relatively high. In many cases, high concentrations like these can interfere with purification, or affect the activity of the protein (Szundi et al., 2021). In those kinds of cases where gentle solubilization of proteins is required, it is common practice to use additives at concentrations as low as possible in order to minimize detrimental effects. DDM and LMNG are typically used from 0.1% to 1%, whereas SMAs have been successfully used in concentrations as low as 0.25% (Kopf et al., 2020; Pitch et al., 2021). In order to determine useful minimal concentrations, we performed a second extraction screen with concentrations of 0.125%, 0.25%, 0.5%, 1%, and 2.5% for a selected panel of polymers and non-ionic detergents. Protein identifications and quantities from this experiment can be found in Table S2. In line with the first screening, DDM and LMNG were the most efficient compounds and reached maximum membrane protein counts at concentrations as low as 0.5% (Figure 5A) and maximum summed intensities at 1% (Figure 5B). In agreement with our earlier experiments, Ultrasolute^TM^ Amphipol-17 and -18 showed very high solubilization efficiency reaching their maximum counts at 1% and plateauing at a maximum value of summed intensities comparable to detergents. AASTY 11-45 and 6-50 also reached a plateau for summed intensities at concentrations above 1%, although only AASTY 11-45 does not seem to find a plateau for the number of individual membrane proteins (Figure 5A,B). DIBMAs in general seem to find a plateau for both quantities, although with much lower values (Figure 5A,B), arguing for a generally lower solubilization efficiency. For all other compounds, the curves imply that the concentration for maximum solubilization has not been reached and may clearly exceed 2.5%. Combined, this shows that polymers can be efficiently used at concentrations significantly lower than 2.5%. We would recommend 0.25 – 1% for Ultrasolute^TM^ Amphipols and 0.5 – 1% for AASTYs, while a higher concentration would be required for DIBMAs and SMAs. To what extent these results from endogenously expressed targets in this study can be transferred to overexpression conditions has to be tested individually for each target.

**Figure 5.**
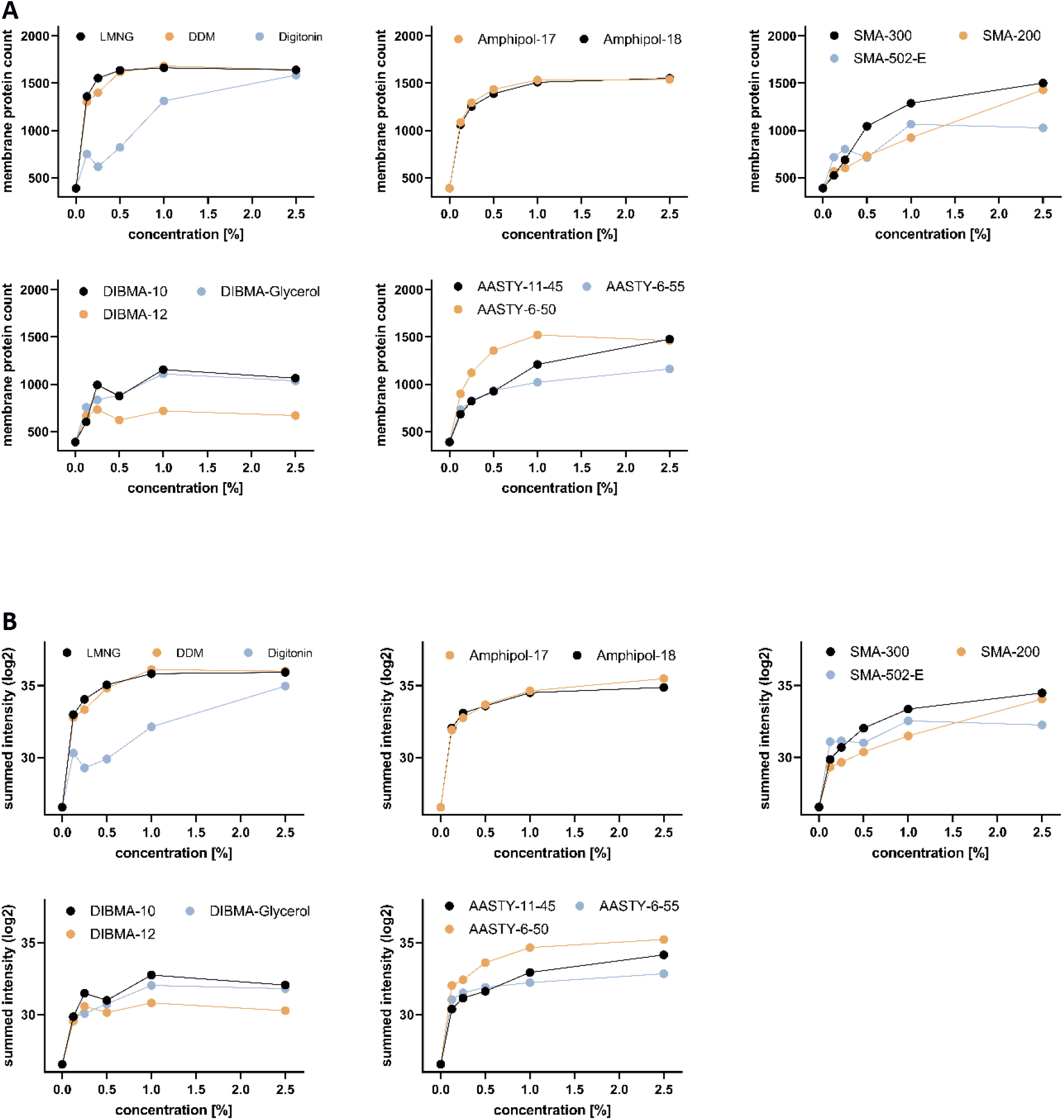
Membrane protein extraction efficiencies of tested solubilization agents. **A.** Number of unique membrane proteins detected as a function of solubilization agent concentration concentrations (0.125 %, 0.25 %, 0.5 %, 1 %, and 2.5 %). **B.** Summed membrane protein intensities as a function of solubilization agent concentration.

Finally, to further dissect the versatility of these polymers we created a heatmap of the relative protein intensities obtained for each individual detected membrane protein, at each different solubilization agent concentration (Figure 6). Unsurprisingly, detergents are able to solubilize most of the proteins in the map with high efficiency. In addition, Ultrasolute^TM^ Amphipols and to a lesser extent AASTY 6-50 produce a very similar pattern, both in terms of concentration dependency and number and identity of the solubilized proteins. Combined this demonstrates that nanodisc-forming polymers present a strong alternative to detergents, especially when the lipid environment is key for protein function.

**Figure 6.**
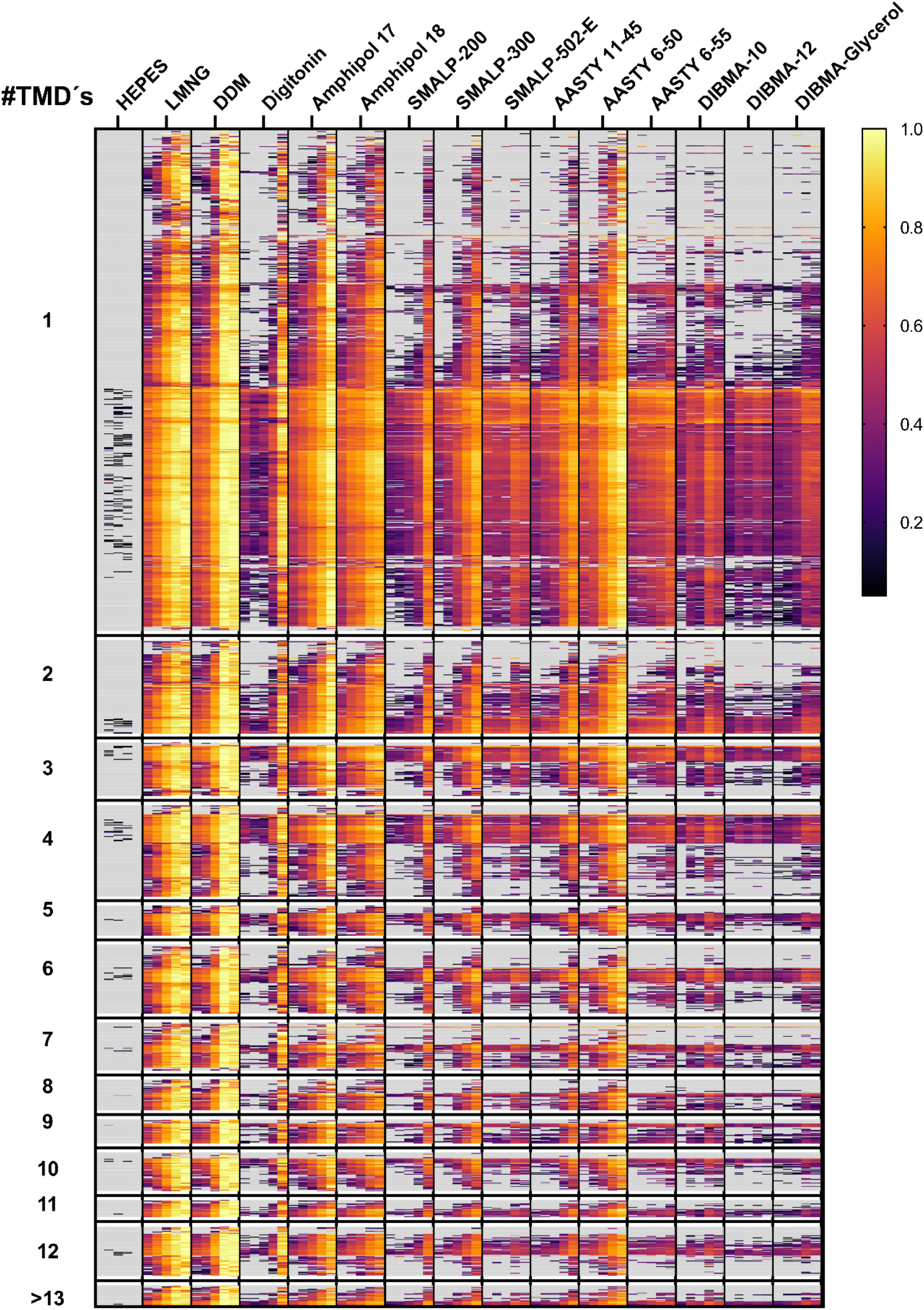
Relative membrane protein intensities in relation to solubilization agent concentrations. For each solubilization agent, five increasing concentrations were tested (0.125 %, 0.25 %, 0.5 %, 1 %, and 2.5 %) and plotted from left to right. HEPES buffer without any additives was used as control in triplicate. Log2 intensities were interval normalised for each row and missing values are displayd in grey. Proteins were grouped according to their number of transmembrane domains as indicated.

## Concluding remarks

We used LC-MS to perform label-free quantification proteomics aimed at screening native-nanodisc-forming polymers and compared them with state-of-the-art detergents for their ability to solubilize the Hek293 membrane proteome. We omitted other membrane-mimetic strategies — such as MSP-based nanodisc (Bayburt et al., 2002), peptidisc (Carlson et al., 2018), and saponins (Frauenfeld et al., 2016) — as they all require the intermediate addition of detergents, thus altering the native environment of the proteins under investigation. The study presented here is the first major attempt to compare the ability of a comprehensive panel of polymers at the membrane proteome scale. As such, we believe that this study will become a reference for researchers working with different aspects of the biochemistry of membrane proteins.

We have demonstrated that these synthetic polymers exhibit solubilization performance that can rival those observed in commonly used detergents, but have the advantage of preserving the protein in a near-native environment; Ultrasolute^TM^ Amphiphol 17 and 18 were as efficient as the denaturing detergent group. Considering the separate clustering observed in Figure 1, plus the patterns observed in Figure 6, we believe that, for any given target, it is likely that at least one polymer class should provide efficient solubilization. We are certain that the ability to preserve the native membrane environment will become and invaluable tool for membrane biochemists. As such, we anticipate that these synthetic polymers will have a broad range of applications in the future, in particular as components of native lysis and solubilization buffers. Moreover, with the ever-increasing access the electron cryo-microscopy we expect that these polymers will make a big impact in the structural biology of integral membrane proteins.

## Acknowledgments

All mass spectrometric analysis was performed in the CMMC/CECAD Proteomics Facility, CECAD University of Cologne.

## Data Availability

The results of the mass spectrometry data included in this manuscript is available in Table S2.

---

**Supplementary Figure 1.**
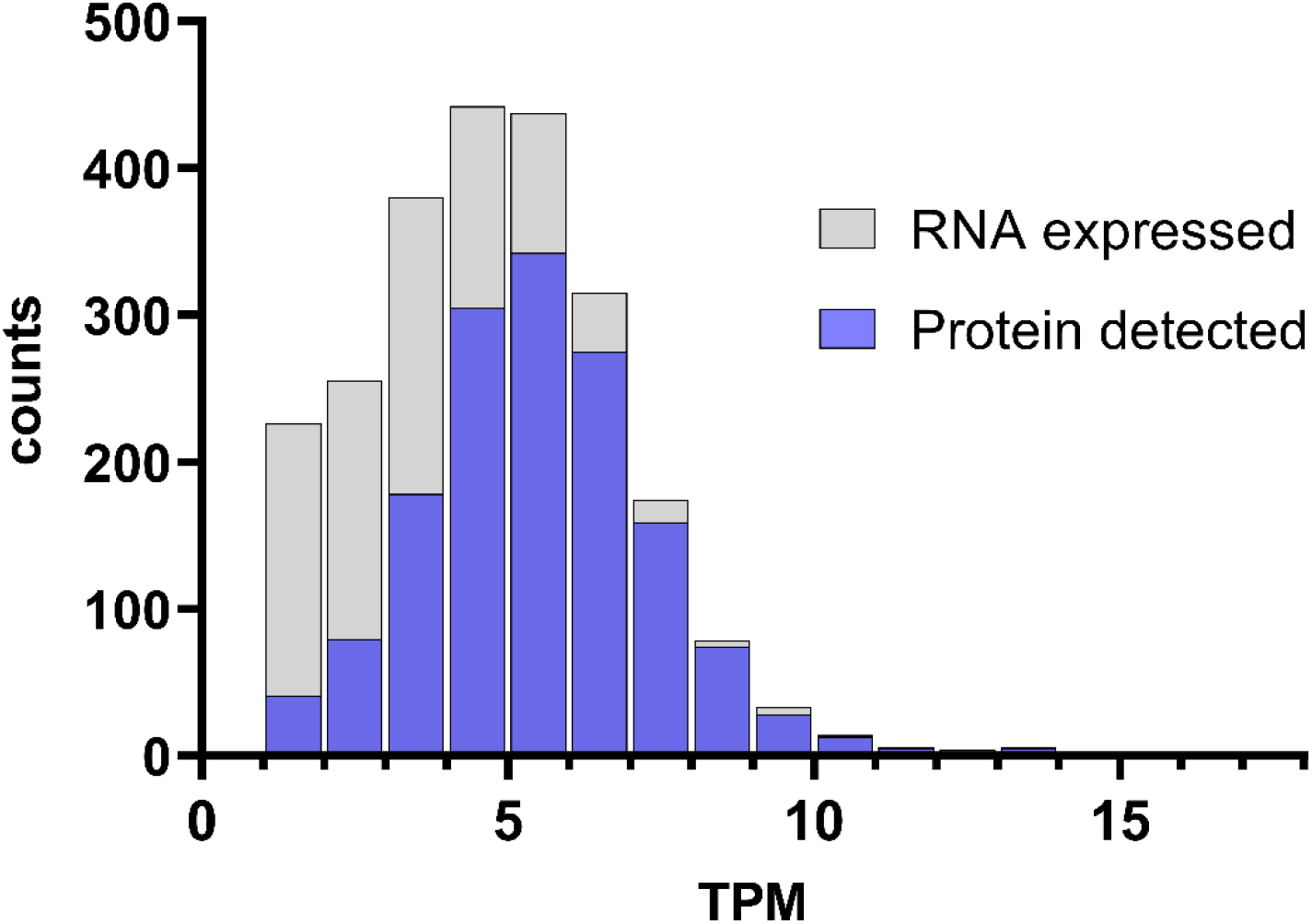
Correspondence between membrane protein mRNA and protein detection in Hek293 cells. RNA transcript counts of membrane protein coding genes were plotted against binned log2 transformed TPM. Transcripts with TPM < 1 were removed. In blue, we show the number of proteins we could detect as a function of their corresponding TPM value obtained from Protein Atlas (See methods).

**Supplementary Table 1.** Physicochemical characteristics of all solubilization agents used.

| Copolymer/detergent | Apolar subunit | Polar subunit | Subunit ratio | CMC [% (w/v)] | Mw [g/mol] | Charge | Source |
| --- | --- | --- | --- | --- | --- | --- | --- |
| AASTY 11-55 | Styrene | Acrylic acid | 55:45 | n.a. | 11,000 | Negative | Cube Biotech |
| AASTY 6-55 | Styrene | Acrylic acid | 55:45 | n.a. | 5,500 | Negative | Cube Biotech |
| AASTY 11-50 | Styrene | Acrylic acid | 1:1 | n.a. | 11,000 | Negative | Cube Biotech |
| AASTY 6-50 | Styrene | Acrylic acid | 1:1 | n.a. | 5,500 | Negative | Cube Biotech |
| AASTY 11-45 | Styrene | Acrylic acid | 45:55 | n.a. | 11,000 | Negative | Cube Biotech |
| Ultrasolute Amphipol-18 (CyclAPol C8C0-50) | Cyclooctyl acrylamide | Acrylic acid | 1:1 | n.a. | 7,500 | Negative | Cube Biotech |
| Ultrasolute Amphipol-17 (CyclAPol C6C2-50) | 2-Cyclohexyl-ethyl acrylamide | Acrylic acid | 1:1 | n.a. | 7,500 | Negative | Cube Biotech |
| DIBMA Glucosamine | Diisobutylidene | Maleamic acid-glucosamine | 1:1 | n.a. | 14,900 | Negative | Cube Biotech |
| DIBMA Glycerol | Diisobutylidene | Maleamic acid-3-amino-1,2-propanediol | 1:1 | n.a. | 12,500 | Negative | Cube Biotech |
| DIBMA 12 | Diisobutylidene | Maleic acid | 1:1 | n.a. | 12,000 | Negative | Cube Biotech |
| DIBMA 10 | Diisobutylidene | Maleic acid | 1:1 | n.a. | 10,000 | Negative | Cube Biotech |
| SMA 502-E | Styrene | Maleic acid monoethyl ester | 1.5:1 | n.a. | 7,000 | Negative | Cube Biotech |
| SMA 300 | Styrene | Maleic acid | 3:1 | n.a. | 10,000 | Negative | Cube Biotech |
| SMA 200 | Styrene | Maleic acid | 2:1 | n.a. | 6,500 | Negative | Cube Biotech |
| Digitonin | n.a | n.a | n.a | 0.2 - 0.3 | 1,229.31 | Neutral | Carl Roth |
| Lauryl maltose neopentyl glycol (LMNG) | n.a | n.a | n.a | 0.001 | 1,005.19 | Neutral | Anatrace |
| Decy-β-maltoside (DM) | n.a | n.a | n.a | 0.087 | 482.56 | Neutral | Cube Biotech |
| Dodecyl-β-maltoside (DDM) | n.a | n.a | n.a | 0.0087 | 510.62 | Neutral | Cube Biotech |
| RIPA buffer | n.a | n.a | n.a | n.a. | n.a. | mixed | Sigma-Aldrich |
| Sodium dodecylsulfat (SDS) | n.a | n.a | n.a | 0.2 | 288.38 | Negative | Sigma-Aldrich |
| Foscholine-12 (Fos-12) | n.a | n.a | n.a | 0.047 | 351.5 | Zwitterionic | Cube Biotech |
| Sodium deoxycholate (SDC) | n.a | n.a | n.a | 0.083-0.325 | 414.55 | anionic | Sigma-Aldrich |

